# The codon frustration index as a new metric for mRNA stability, translation efficiency, and rates of protein synthesis

**DOI:** 10.1101/692996

**Authors:** Sergio Forcelloni, Andrea Giansanti

**Affiliations:** Sapienza University of Rome, Department of Physics, P.le A. Moro 5, 00185 Roma, Italy; Istituto Nazionale di Fisica Nucleare, INFN, Roma1 section. 00185, Roma, Italy

**Keywords:** codon usage bias, tRNA, frustrated codon, translation efficiency, translation accuracy, mRNA stability

## Abstract

Taking the human genome as a case of study, we propose a new classification of codons based only on two genomic information. We use the relative synonymous codon usage (RSCU) as a measure of non-uniform usage of synonymous codons. Similarly, we introduce here the *relative gene frequencies of cognate tRNAs (RGFCt)* to quantify the non-uniform availability of cognate tRNAs in each family of synonymous codons. Using these two quantities, we define two general groups of codons: *non-frustrated codons*, whose usage in the coding sequences is in proportion to the expected cognate tRNA levels, and *frustrated codons*, which do not satisfy this proportionality. With this decoding for every codon, we defined the *Codon Frustration Index (CFI)* as the net frustration of a gene, normalized for its length. Notably, we find that CFI correlates very well with other independent measures of CUB and a high content of non-frustrated codons increases both translation efficiency and mRNA stability. Finally, we show that genes with either a high content of frustrated or of non-frustrated codons are differentially enriched in specific functional classes that typically comprise nucleic acid binding proteins, mRNA processing factors, RNA helicase, and in several transcription factors.

## 1 Introduction

The genetic code is redundant, with most amino acids encoded by several synonymous codons. The usage of such synonymous codons in protein coding sequences is generally not random, a phenomenon called codon usage bias (CUB) [1–5]. The prevailing hypothesis to explain the biological significance of CUB revolves around the concept that certain codons are translated both faster and more accurately [6]. Although several others factors may also contribute [7–10], the occurrence of synonymous codons and the availability of tRNAs capable of decoding them are considered the major factors that govern translation efficiency and accuracy [5,6,11–13]. Thus, codons that are recognized by abundant or rare tRNAs have been respectively referred to as fast and slow codons (or as codons that are highly or poorly adapted to the intracellular tRNA pool) [14].

During translation, specific tRNAs recognize the codons in the mRNA and deliver the corresponding amino acids to the ribosome. Every organism can express several isoacceptor tRNAs that translate synonymous codons for the same amino acid. The concentrations of tRNAs vary markedly in the cellular pool, and it determines the codon translation efficiency in both unicellular and multicellular organisms [2;15–18], including human [19,20]. Because mature tRNAs in humans are thought to be very stable, tRNA levels are mostly determined by tRNA transcription rates [21]. Unfortunately, accurate prediction and experimental determination of the intracellular levels of tRNA are technically challenging [22]. The genomic tRNA content has been quite used as a proxy for the tRNA cellular abundance [14,16–20,23–27]. Therefore, the tRNA gene copy number determines relative concentrations of isoacceptor tRNAs that, in turn, determine codon translation efficiency [28].

Considering different eukaryotes, Qian et al. found that the optimal strategy in enhancing translation efficiency consists in using codons in proportion to cognate tRNA concentrations (*proportional rule*) [29]. Accordingly, if codons are used in proportion to the availability of the cognate tRNAs, then this could dampen any pausing effect since rare codons matching rare tRNAs will not be as rate-limiting as if they were used more often [8].

In this study, we propose a new classification of codons that reflects the proportional rule by Qian et al. [29], and it is based only on statistical information in the human karyotype. We used the relative synonymous codon usage (RSCU) as a measure of non-uniform usage of synonymous codons in the protein coding sequences. Similarly, we introduced the *Relative Gene Frequency of Cognate tRNAs (RGFCt)*, considering the tRNA gene copy numbers as a rough proxy of intracellular tRNA levels. Using these two definitions, we distinguished between: i) *non-frustrated codons*, whose usage in the coding sequences is in proportion to the expected level of the cognate tRNAs in the cell, and ii) *frustrated codons*, which do not satisfy this proportionality.

We also defined the *codon frustration index (CFI)* as the average of frustrations of the codons in each gene. A strong correlation was observed between the CFI and independent measures of CUB such as tAI [17], CompAI [30], and *N_c_* [31]. In line with the proportional rule by Qian et al. [29], we found that a high content of non-frustrated codons increases the translation elongation rates. Moreover, we detected a strong positive correlation between CFI and mRNA stability. Finally, we evaluated the statistical significance of this new metric. We found that genes with a high content of non-frustrated codons and genes with a high content of frustrated codons are significantly enriched in non-overlapping functional protein classes, related to nucleic acid binding proteins, mRNA processing factors, RNA helicase, and in many types of transcription factors.

## 2 Materials and Methods

### 2.1 Data sources

The human proteome was downloaded from the UniProtKB/SwissProt database (http://www.uniprot.org/uniprot) [32]. We selected human proteins by searching for manually annotated and reviewed proteins belonging to the organism Homo Sapiens (Organism ID: 9606, Proteome ID: UP000005640). Human protein coding sequences were retrieved by Ensembl Genome Browser 94 [33]. We included only coding sequences that start with AUG, end with a stop codon (UAG, UAA, or UGA), have a length that is a multiple of three, and do not have unidentified bases. Each coding sequence was translated in the corresponding amino acid sequence and all sequences that do not have a complete correspondence with a protein sequence in UniProtKB/SwissProt were removed from the analysis. The resulting dataset contained 18150 human coding sequences. Human tRNA gene copy numbers (tGCN) were retrieved from the Genomic tRNA Database (GtRNAdb 2.0) (http://lowelab.ucsc.edu/GtRNAdb/Hsapi) [34].

### 2.2 Relative synonymous codon usage

The relative synonymous codon usage (RSCU) is the observed frequency of a synonymous codon divided by the expected frequency if all the synonymous codons for the amino acid were used uniformly [35]. Formally, the RSCU of codon *j* encoding for the amino acid *i* (*RSCU_ij_*) is defined as:

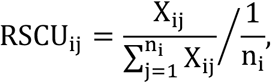

where *n_i_* is the number of synonymous codons encoding for the amino acid *i, X_ij_* is the number of occurrences of the codon *j*, and the summation is taken over all synonymous codons. If the RSCU is close to 1, synonymous codons are used without apparent biases. RSCU values greater or less than 1 indicate that the corresponding codons are used more or less frequently than expected, respectively.

### Relative gene frequency of cognate tRNAs

In the human genome as well as in all other eukaryotes there is a redundancy in the set of tRNAs genes. The number of tRNAs genes in the human genome (also referred to as the tRNA gene copy number - tGCN) associated with the 20 amino acids varies from 7 (Trp) to 34 (Ala) (see http://lowelab.ucsc.edu/GtRNAdb/Hsapi). The tGCN also varies internally to the same codon family with some codon associated with a higher or lower number of tRNA genes than other synonymous codons. In analogy with the definition of the RSCU, we introduce here the *Relative Gene Frequency of Cognate tRNAs (RGFCt)*, which is basically a reassessment of the Relative Gene Frequency (RGF) [16] for taking into account the wobble rules [36]. In particular, the protocol to calculate the RGFCt of a codon *j* encoding for the amino acid *i* (*RGFCt_ij_*) is as follow: a) we estimate the total gene copy number of tRNAs that serve in translating it, incorporating tRNAs that contribute to translation through both Watson-Crick (WC) and wobble (wb) pairing rules (*cognate tRNA gene copy number*); b) we extrapolate all other synonymous codons for the amino acid *i* and all the corresponding tRNA anticodons satisfying WC or wb pairing rules; c) we compute the average of the cognate tRNA gene copy numbers associated with such family of synonymous codons; d) finally, the formula used for estimating the *RGFCt_ij_* is:

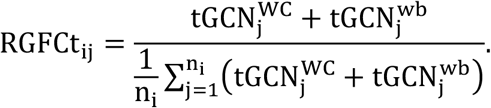

Where 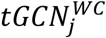 and 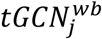 are the gene copy number of tRNAs with WC and wb complementarity with codon *j*, respectively; n_i_ is the number of synonymous codons encoding for amino acid *i*; the summation is taken over all synonymous codons. If the *RGFCt* = 1, then the cognate tRNA gene copy number for that synonymous codon is equal to the average one; conversely, if the *RGFCt* is > 1 or < 1, then the cognate tRNA gene copy number for that synonymous codon is greater or less than the average one, respectively.

In this study, we do not consider tissue-specific differences in tRNA levels since their experimental determination remains elusive and unavailable on a broad scope [3,22]. Thus, and in line with previous studies in human [19,20,23–26], we used the tRNA gene copy number as a first approximation for the intracellular tRNA level (a similar approach is also adopted in the definition of tAI [17] and CompAI [30]).

### RSCU-RGFCt plane and definition of frustrated and non-frustrated codons

In the *RSCU-RGFCt plane*, the RSCU value is plotted as the ordinate, and the RGFCt value is plotted as the abscissa. In Fig. 1, we report the RSCU-RGFCt plane for the human genome.

**Figure 1:**
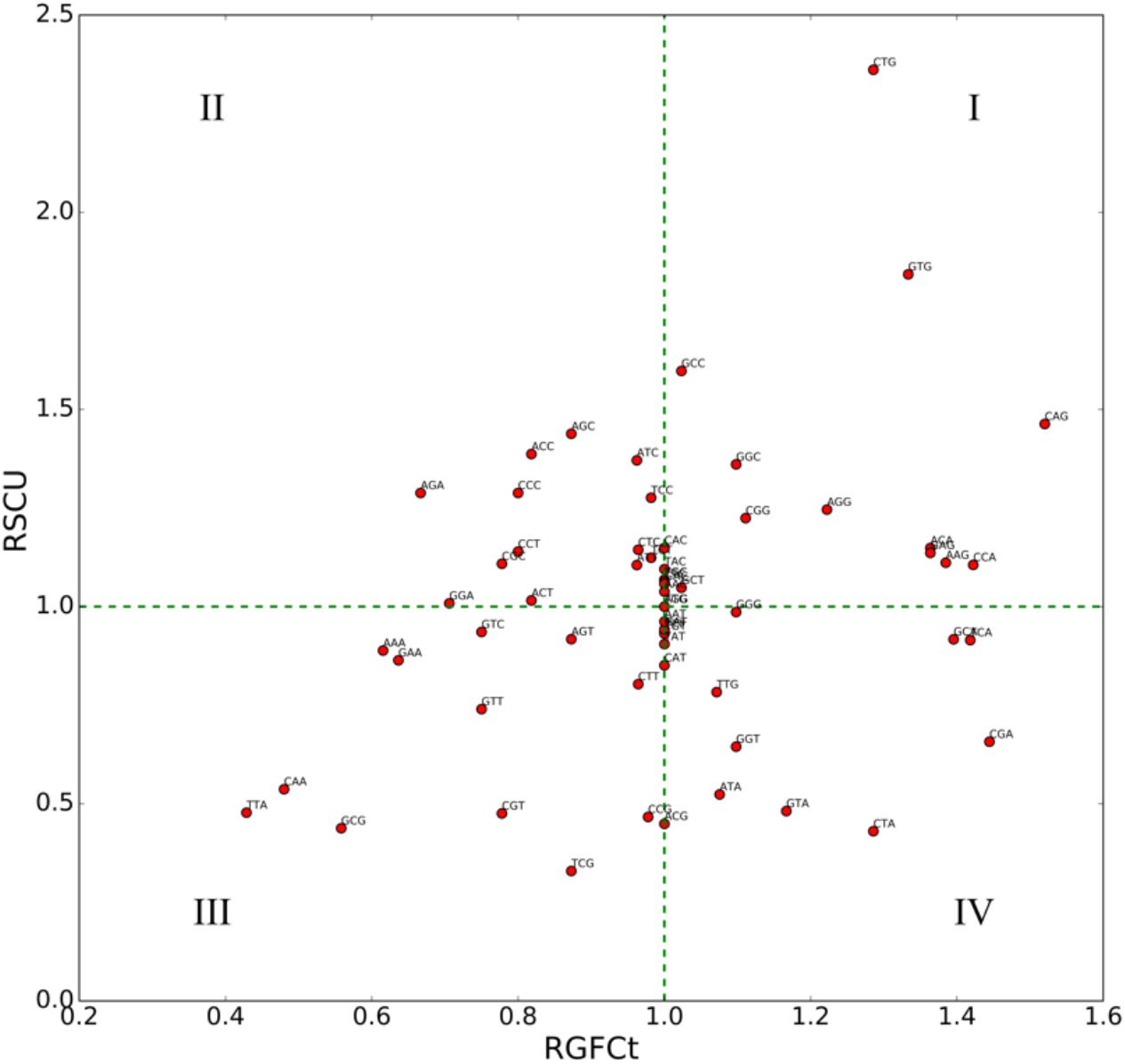
The RSCU-RGFCt plane, and the human genome. Each of the 61 codons is reported as a single point on this plane. The two broken lines represent the exceptional cases *RSCU* = 1 and *RGFIt* = 1. Codons in the I and III quadrants are referred to as non-frustrated, and codons in the II and IV quadrants as frustrated.

Considering the two broken lines corresponding to the exceptional cases *RSCU* = 1 and *RGFCt* = 1, we can distinguish between codons that are used proportionally to the genomic content of the corresponding cognate tRNA and codons that do not satisfy this proportionality. We denoted codons in the II and IV quadrants as *frustrated* because they correspond to *high/low* RSCU values but *low/high* RGFCt values, and codons in the I and III quadrants as *non-frustrated* because they correspond to *high/low* RSCU values, and coherently *high/low* RGFCt values. Notably, 14 out of 17 optimal codons detected by Comeron [24] are classified as non-frustrated in this metric. For how the codon usage patterns and tRNA gene copy numbers have coevolved, 13 codons, other than of those encoding for Met and Trp, are located on the vertical line *RGFCt* = 1. To classify these codons and decide which of the two groups they belong to, we used data about the mean typical decoding rate (MTDR) of individual codons provided by Dana and Tuller [37]. In Fig. S1, we report a boxplot of MTDR for codons with *RSCU* > 1 and codons with *RSCU* < 1. Codons with *RSCU* > 1 are read faster than codons with *RSCU* < 1 in a statistically significant way (*p* – *value* < 0.05). Therefore, we associated codons with *RSCU* > 1 and *RGFCt* = 1 to the non-frustrated group and codons with *RSCU* < 1 and *RGFCt* = 1 to the frustrated group.

In analogy with the Ising model in statistical physics [38], we assigned spin up (*s* = 1) to non-frustrated codons and spin down (*s* = −1) to frustrated codons. Then, we translated each coding sequence into a sequence of spin variables and defined the *codon frustration index (CFI)* as the average frustration of the gene’s codons:

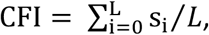

where *L* is the length of the coding sequence; *s_i_* is the spin variable associated with the codon in position *i*. Non-degenerate codons (Met and Trp) were discarded from the calculation. If the CFI is close to 0, frustrated and non-frustrated codons are equally used without apparent biases. CFI values greater or less than 0 indicate that non-frustrated or frustrated codons are favored in the coding sequence, respectively.

### Codon bias indices

In this work, we used different independent measure for quantifying CUB of individual genes: the *tRNA adaptation index* (tAI) [17], which estimates the extent of adaptation of a gene to its genomic tRNA pool; the *competition adaptation index* (CompAI) [30], which is based on the competition of cognate and near-cognate tRNAs to bind to the A-site on the ribosome during translation; the *effective number of codons* (*N_c_*) [31], which is a statistical measure of the number of different codons used in a coding sequence.

### Prediction of the translational rates

The mean typical decoding rates (MTDR) index [37] was used to estimate the translation elongation rate of a gene. This index is based on the typical codon decoding times from Ribo-seq data after filtering biases and phenomena such as ribosomal traffic jams and translational pauses.

### Prediction of the mRNA stability

The mRNA stability was assessed as the arithmetic mean of the codon stabilization coefficient (CSC) of the gene’s codons [39]. The CSC is defined as the Pearson correlation coefficient between mRNA half-life and codon occurrence. mRNAs enriched in codons displaying a positive correlation tend to be more stable; conversely, mRNAs enriched in codons showing a negative correlation tend to be less stable. In this study, we used the CSC values for each codon derived by Wu et al. [39] for four different human cells (HEK293T, Hela, K562, RPE).

### Shuffling algorithm of protein coding sequences

The statistical significance of the CFI, as a measure of an evolutionary signal against a random case, was assessed through a shuffling procedure of the coding sequences composed by four steps [40]. Firstly, we count the occurrences of each codon in the whole dataset of coding sequences. Secondly, we arranged codons in families according to the amino acid they translate. Thirdly, we calculate the frequency of each codon within its synonymous family. Finally, we re-read each coding sequence and replace each codon with a codon in the same family, by selecting it randomly and proportionally to its frequency calculated in the third step. Thus, this algorithm replaces codons in each coding sequence while preserving the amino acid sequence. For each coding sequence, we repeat the shuffling procedure 1000 times, re-calculating for each sample the CFI value. Thus, we associate to each gene the distribution of CFIs generated by the null model of random codon choice. We then associate to the effective CFI of each gene (abscissa of figure 4) the corresponding Z-score evaluated as

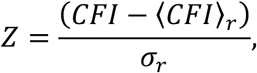

where 〈*CFI*〉_*r*_ and *σ_r_* are the average value and the standard deviation of the distribution of CFIs generated by the null model, respectively. Values of CFI which exceed the 95% confidence level band (i.e., −1.96 and +1.96 standard deviations) are considered as statistically significant, corresponding to events that are hardly sampled (*p* − value < 0.05) in the null, random model.

### Functional classification

The functional analyses were performed using PANTHER (Protein Annotation Through Evolutionary Relationship) classification system (http://www.pantherdb.org) [41,42].

## Results

### Codon frustration index and other measures of codon usage bias

We first studied the correlation between the CFI for individual genes and other independent measures of CUB such as tAI, CompAI, MTDR, and *N_c_* (Fig. 2).

**Figure 2:**
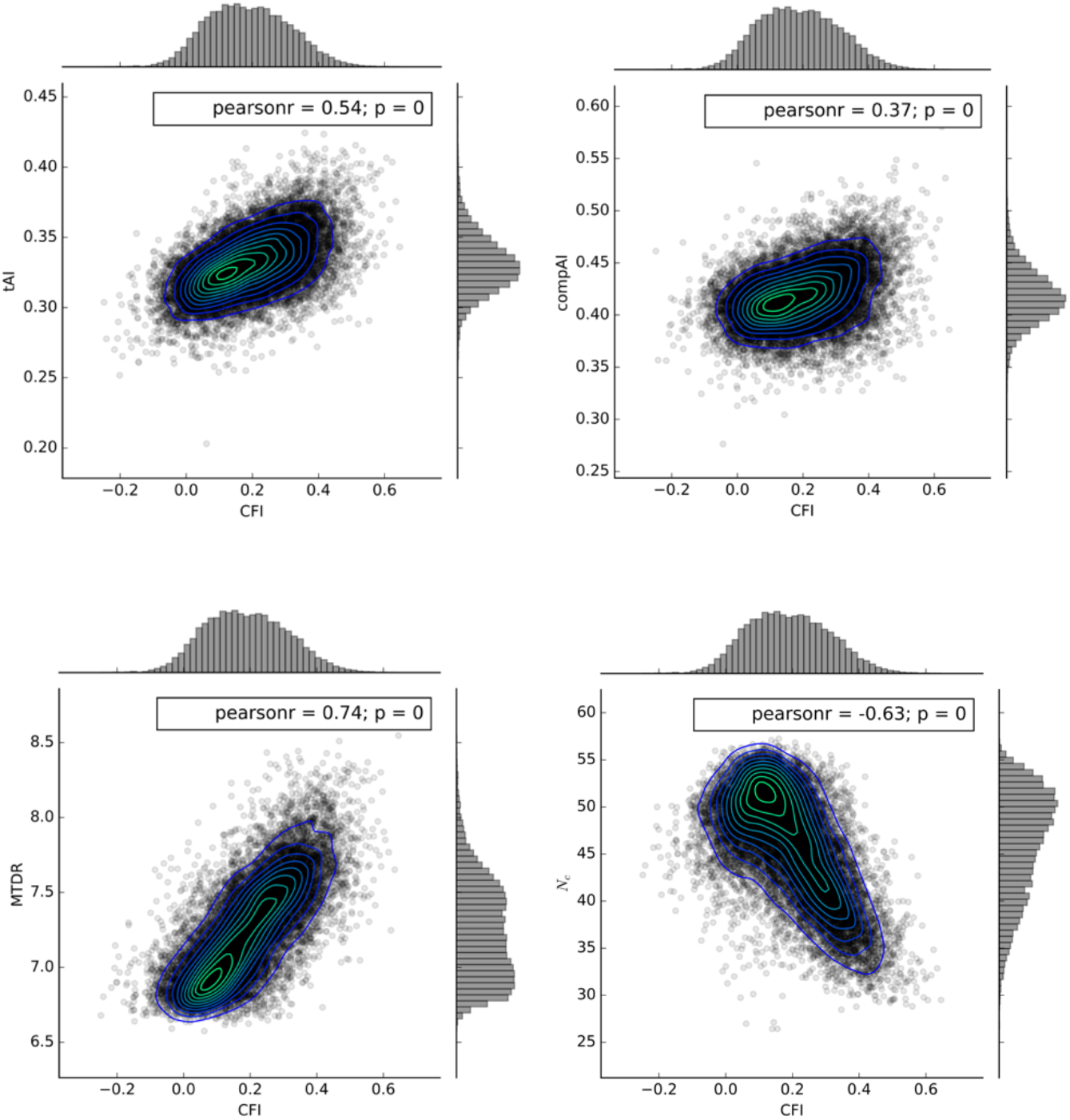
Correlation between CFI and other measures of CUB (i.e., tAI, CompAI, MTDR, and *N_c_*). Each point represents a single gene. Pearson’s correlation coefficients and the corresponding p-value are reported at the top of each panel.

The CFI is positively correlated with MTDR (r=0.74), tAI (r=0.54), CompAI (r=0.37), and negatively with *N_c_* (r=−0.63). Notably, the strong positive correlation between CFI and MTDR means that a preferential usage of non-frustrated codons increases the translation elongation rate. This is coherent with the results by Qian et al. [29], who stated that the optimal usage of synonymous codons for maximizing translational efficiency is to use them in proportion to the availability of cognate tRNAs in the cell.

### Codon frustration index and mRNA stability

mRNA codon composition is emerging as an essential factor in determining mRNA stability and translation efficiency [5,39,40]. To investigate how mRNA stability relates to the CFI of individual genes, we used the codon stabilization coefficient (CSC) metric, which ranks codons according to their contribution to mRNA stability. We used CSC values, derived by Wu et al. [39], for four human cell lines (HEK293T, Hela, K562, RPE), and calculated the arithmetic mean of CSC for each gene (CSCg). Genes enriched in codons with a negative CSC value are less stable; conversely, genes enriched in codons with a positive CSC value are more stable. In Fig. 3, we show the correlations between CFI and CSCg for individual genes.

**Figure 3:**
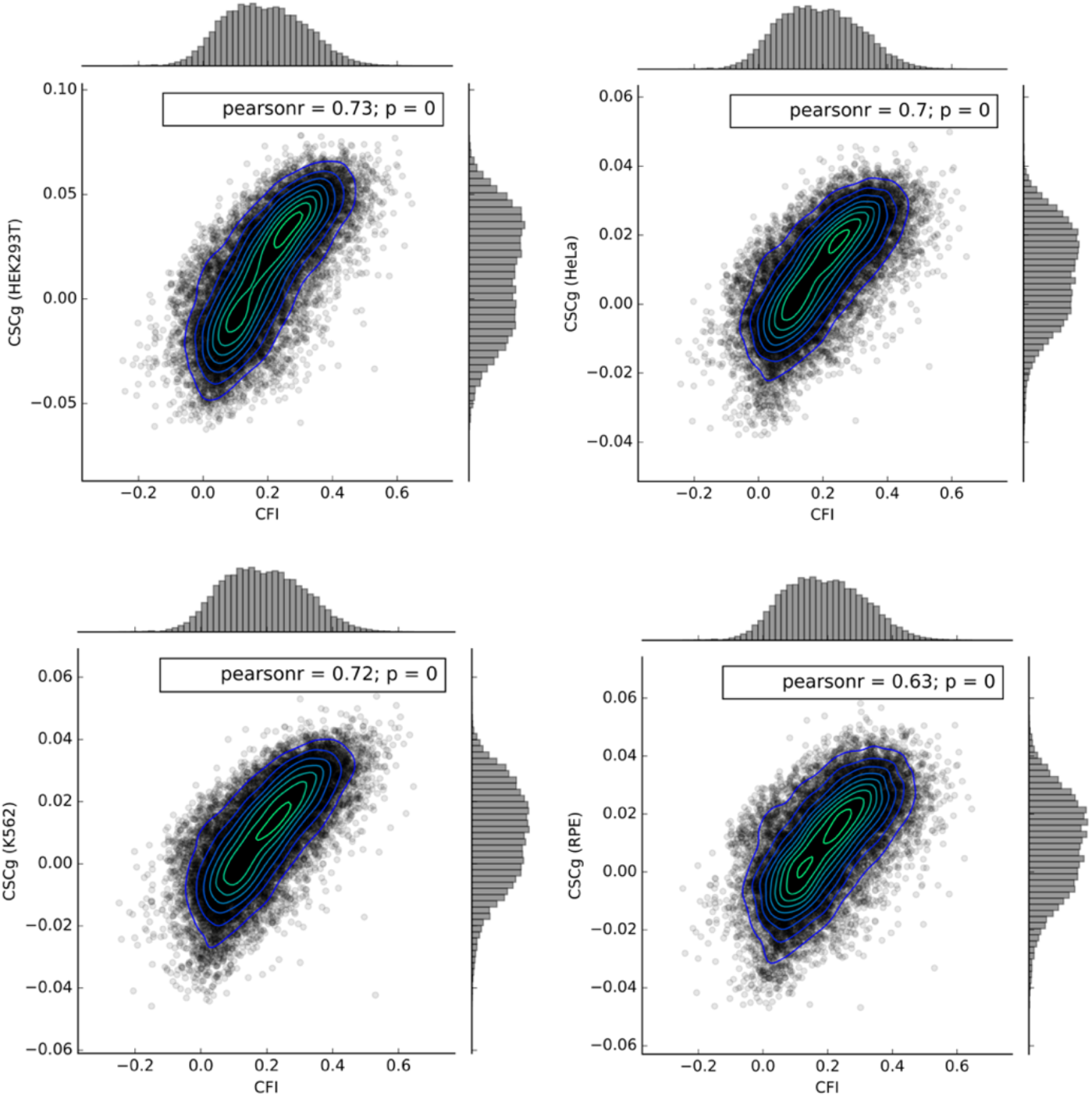
Correlations between CFI and CSCg for four human cell lines (HEK293T, Hela, K562, RPE). Each point represents a single gene. Correlations are calculated using CSC values for individual codons by Wu et al. [39]. Pearson’s correlation coefficients and the corresponding p-values are shown at the top left of each panel.

CFI is strongly and positively correlated with the CSCg for individual genes in all human cell lines. Accordingly and in line with the fact that a preferential usage of non-frustrated codons tends to increase translation efficiency (Fig. 2), high content of non-frustrated codons also tends to enhance mRNA stability.

### Functional classification of genes with a significant codon frustration index

A codon shuffling algorithm was used to test the statistical significance of the CFI values against the null model made of random patterns of synonymous codons (see Materials and Methods for details). In Fig. 4, we show the CFI of individual genes on the abscissa and the corresponding Z-score on the ordinate.

**Figure 4:**
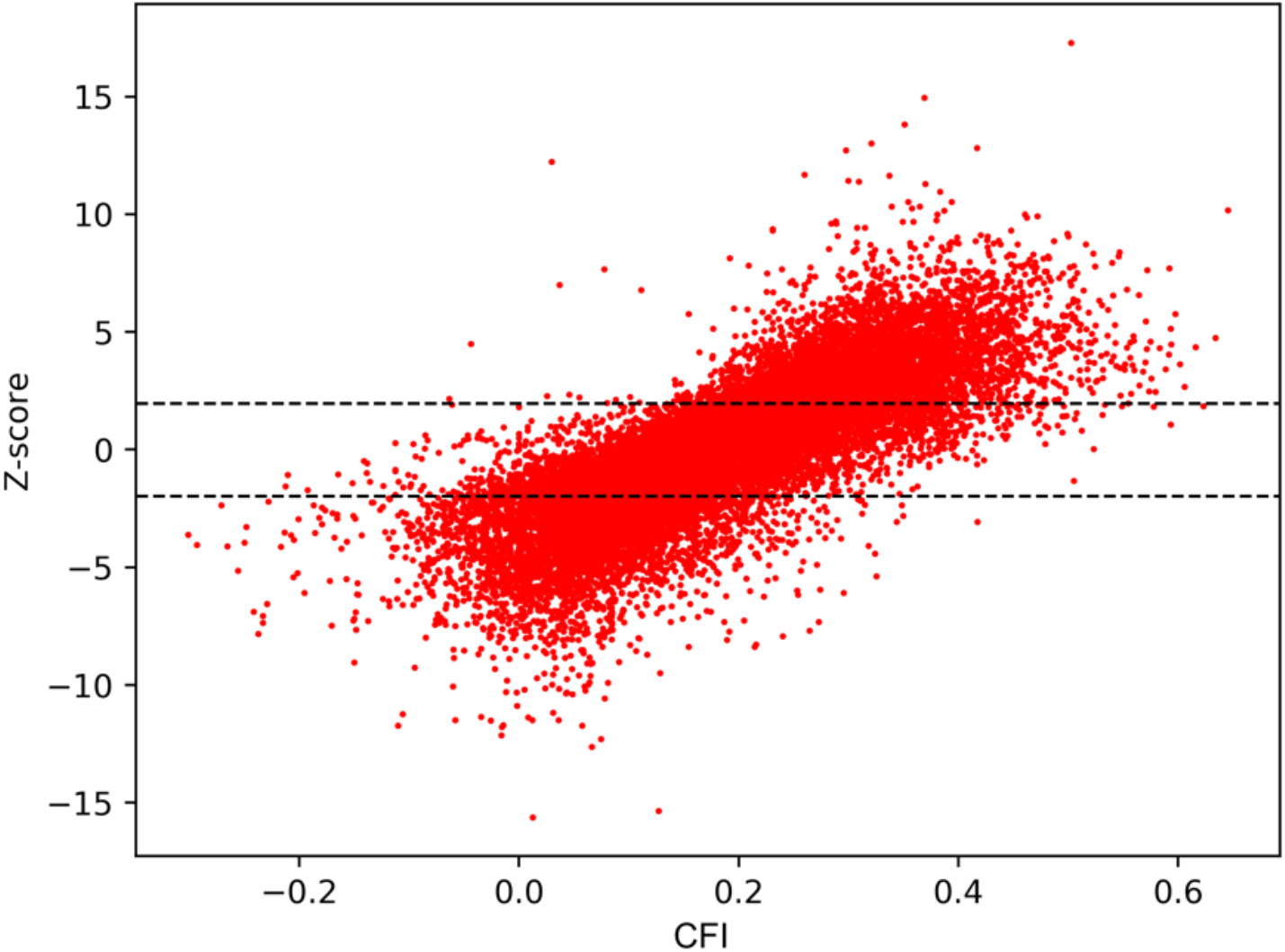
CFI of individual genes (abscissa) vs. Z-score relative to what expected in the random case (ordinate). Black dotted lines identify the critical Z score values for a 95% confidence level (i.e., −1.96 and +1.96 standard deviations). If the Z score falls outside −1.96 and +1.96, the corresponding CFI value was considered as statistically significant (*p* − *value* < 0.05).

A PANTHER functional classification was performed separately for the 5026 genes having a *Z* − score > 1.96 (Table 1), and for the 4558 genes having a *Z* − *score* < −1.96 (Table 2).

**Table 1:**
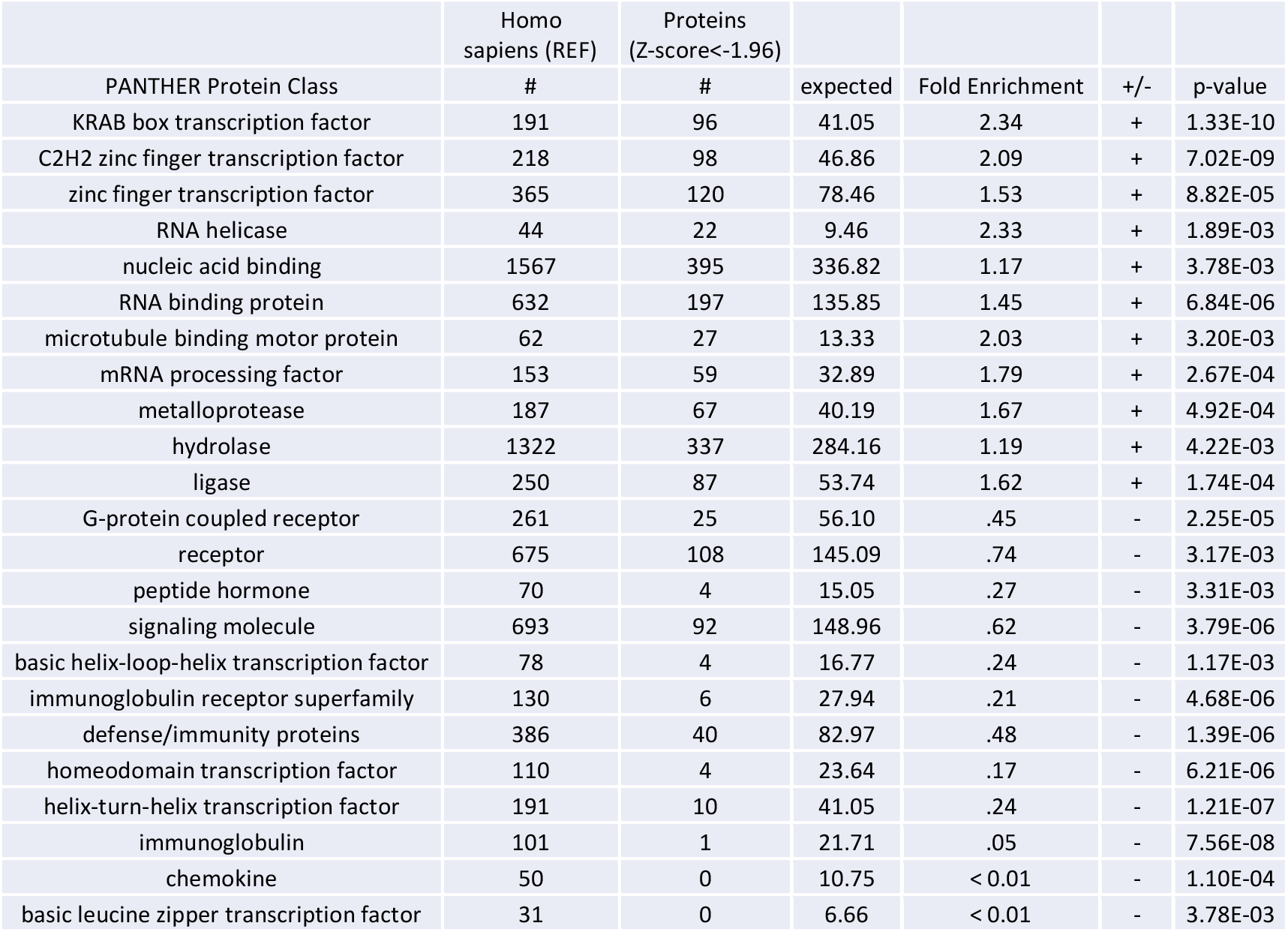
Fold enrichment of genes with a standard score lower than −1.96 in PANTHER protein classes. Only statistically significant differences are reported (*p* − *value* < 0.05).

|  | Homo sapiens (REF) | Proteins (Z-score<-1.96) |  |  |  |  |
| --- | --- | --- | --- | --- | --- | --- |
| PANTHER Protein Class | # | # | expected | Fold Enrichment | +/- | p-value |
| KRAB box transcription factor | 191 | 96 | 41.05 | 2.34 | + | 1.33E-10 |
| C2H2 zinc finger transcription factor | 218 | 98 | 46.86 | 2.09 | + | 7.02E-09 |
| zinc finger transcription factor | 365 | 120 | 78.46 | 1.53 | + | 8.82E-05 |
| RNA helicase | 44 | 22 | 9.46 | 2.33 | + | 1.89E-03 |
| nucleic acid binding | 1567 | 395 | 336.82 | 1.17 | + | 3.78E-03 |
| RNA binding protein | 632 | 197 | 135.85 | 1.45 | + | 6.84E-06 |
| microtubule binding motor protein | 62 | 27 | 13.33 | 2.03 | + | 3.20E-03 |
| mRNA processing factor | 153 | 59 | 32.89 | 1.79 | + | 2.67E-04 |
| metalloprotease | 187 | 67 | 40.19 | 1.67 | + | 4.92E-04 |
| hydrolase | 1322 | 337 | 284.16 | 1.19 | + | 4.22E-03 |
| ligase | 250 | 87 | 53.74 | 1.62 | + | 1.74E-04 |
| G-protein coupled receptor | 261 | 25 | 56.10 | .45 | - | 2.25E-05 |
| receptor | 675 | 108 | 145.09 | .74 | - | 3.17E-03 |
| peptide hormone | 70 | 4 | 15.05 | .27 | - | 3.31E-03 |
| signaling molecule | 693 | 92 | 148.96 | .62 | - | 3.79E-06 |
| basic helix-loop-helix transcription factor | 78 | 4 | 16.77 | .24 | - | 1.17E-03 |
| immunoglobulin receptor superfamily | 130 | 6 | 27.94 | .21 | - | 4.68E-06 |
| defense/immunity proteins | 386 | 40 | 82.97 | .48 | - | 1.39E-06 |
| homeodomain transcription factor | 110 | 4 | 23.64 | .17 | - | 6.21E-06 |
| helix-turn-helix transcription factor | 191 | 10 | 41.05 | .24 | - | 1.21E-07 |
| immunoglobulin | 101 | 1 | 21.71 | .05 | - | 7.56E-08 |
| chemokine | 50 | 0 | 10.75 | < 0.01 | - | 1.10E-04 |
| basic leucine zipper transcription factor | 31 | 0 | 6.66 | < 0.01 | - | 3.78E-03 |

**Table 2:** Fold enrichment of genes with a standard score higher than 1.96 in PANTHER protein classes. Only statistically significant differences are reported (*p* − *value* < 0.05).

|  | Homo sapiens (REF) | Proteins (Z-score>1.96) |  |  |  |  |
| --- | --- | --- | --- | --- | --- | --- |
| PANTHER Protein Class | # | # | expected | Fold Enrichment | +/- | p-value |
| homeodomain transcription factor | 110 | 63 | 26.05 | 2.42 | + | 9.38E-08 |
| helix-turn-helix transcription factor | 191 | 98 | 45.24 | 2.17 | + | 2.64E-09 |
| transcription factor | 1138 | 377 | 269.54 | 1.40 | + | 1.47E-08 |
| basic helix-loop-helix transcription factor | 78 | 42 | 18.47 | 2.27 | + | 3.75E-05 |
| G-protein coupled receptor | 261 | 135 | 61.82 | 2.18 | + | 1.67E-12 |
| receptor | 675 | 267 | 159.88 | 1.67 | + | 3.46E-12 |
| extracellular matrix protein | 190 | 71 | 45.00 | 1.58 | + | 1.49E-03 |
| DNA binding protein | 457 | 145 | 108.24 | 1.34 | + | 2.35E-03 |
| RNA binding protein | 632 | 103 | 149.69 | .69 | - | 2.44E-04 |
| immunoglobulin receptor superfamily | 130 | 9 | 30.79 | .29 | - | 3.20E-05 |
| defense/immunity protein | 386 | 48 | 91.43 | .53 | - | 6.30E-06 |
| immunoglobulin | 101 | 1 | 23.92 | .04 | - | 1.63E-08 |
| chemokine | 50 | 0 | 11.84 | < 0.01 | - | 3.81E-05 |
| cytokine | 117 | 10 | 27.71 | .36 | - | 6.12E-04 |

Genes with a non-random high content of non-frustrated codons (*Z* − *score* > 1.96) are enriched in receptor, extracellular matrix protein, DNA binding protein, G-protein coupled receptor, and in several transcription factors (i.e., homeodomain, helix-turn-helix, and helix-loop-helix). Conversely, genes with a non-random high content of frustrated codons (*Z* − *score* < −1.96) are enriched in functional classes related to RNA helicase, nucleic acid binding protein, RNA binding protein, microtubule binding motor protein, mRNA processing factor, metalloprotease, hydrolase, ligase, and transcription factors (i.e., KRAB box, and zinc finger). In a nutshell, it is worth noting that genes with a non-random high content of non-frustrated codons (*Z* − *score* > 1.96) are enriched in PANTHER protein classes where genes with a non-random high content of frustrated codons (*Z* − *score* < 1.96) are depleted, and vice versa.

## Discussion

In this work, we propose a new classification of codons that reflects the proportional rule by Qian et al. [29], and it is based only on two statistical indexes extracted from the human genome. The first is the relative synonymous codon usage (RSCU), which is a measure of codon usage bias within each family of synonymous codons. The second is the relative gene frequency of cognate tRNAs (RGFCt), which quantifies the non-uniform availability of cognate tRNAs associated with a family of synonymous codons.

Using these two measures, we introduced the RSCU-RGFCt plane, which provides a species-specific representation of the 61 codons encoding for amino acids. Based on this, we propose a new classification of codons in two groups: i) *non-frustrated codons*, which are used proportionally to the expected level of cognate tRNAs, and ii) *frustrated codons*, which do not follow such proportion.

Compared to the other indices that include tGCNs in their definition (e.g., RGF, tAI, and CompAI), the RGFCt has two main advantages. Firstly, it is a generalization of the classical RGF [16] that takes into account wobble pairing rules [36]. Secondly, it brings out a threshold (*RGFCt* = 1) that permits to classify codons unambiguously according to the number of cognate tRNA genes associated with them.

In analogy with the Ising model in statistical physics [38], we assign spin up (*spin* = +1) to non-frustrated codons and spin down (*spin* = −1) to frustrated codons. Using this coarse-grained grading of each codon we defined the *codon frustration index* (CFI) as the net frustration of the gene, normalized by its length. We found that CFI correlates very well with other independent measures of CUB, such as tAI, CompAI, and *N_c_*. Moreover, a strong positive correlation of CFI was observed, in each gene, with MTDR [37] (which is an effective estimate of the translation elongation rate) and with CSCg [39], which is a proxy for mRNA stability. Thus, we conclude that a high content of non-frustrated codons should be associated to high mRNA stability and translation efficiency. Finally, we found that genes rich in non-frustrated codons and genes rich in frustrated codons have different functional repertoires, suggesting a consistent biological significance of the codon frustration introduced in this study.

## Conclusions

Overall, our results suggest that the distinction between frustrated and non-frustrated codons could represent a new layer of genetic information, rooted in mRNA stability and able to modulate translation efficiency and rates of protein synthesis. It would be tempting to think that in vivo the genetic layer of information we have evidenced here could interfere with the extra control (possibly rooted in epigenetics) represented by the actual in vivo dynamic levels of expressed tRNAs, which vary, in a tissue-specific way, during the cell cycle, and in different conditions associated to development, disease, and stress [43]. When extensive dynamic data on in vivo tRNA levels will be available, it would be very interesting to compare the static picture of the RSCU-RGFCt plane presented here with a dynamical one. In this view, the RSCU-RGFCt plane could provide an integrative representation of the tissue-specific and cell-state-specific balance between the codon usage, resulting from genome evolution, and tRNA levels, which can change dynamically in response to diverse cellular and environmental conditions.

