## Supplementary material for "The codon frustration index as a new metric for mRNA stability, translation efficiency, and rates of protein synthesis": In Fig. S1

**Supplementary Information**

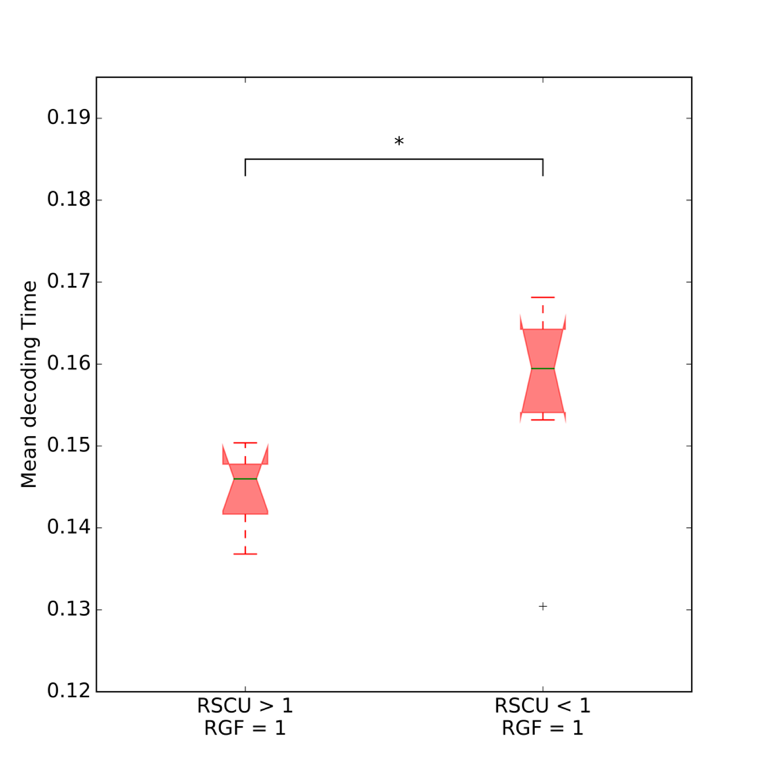

**Figure S1**: Boxplots of the mean decoding time of codons having RSCU > 1 and RGFIt = 1 and codons having RSCU < 1 and RGFIt = 1. The two distributions are statistically different (p value < 0.05).
